# Long-Read Sequencing of the Zebrafish Genome Reorganizes Genomic Architecture

**DOI:** 10.1101/2021.08.27.457855

**Authors:** Yelena Chernyavskaya, Xiaofei Zhang, Jinze Liu, Jessica S. Blackburn

**Affiliations:** Department of Cellular & Molecular Biochemistry, University of Kentucky, Lexington, KY, 40536 USA; Markey Cancer Center at the University of Kentucky, Lexington, KY, 40536, USA; Department of Computer Science, University of Kentucky, Lexington, KY, 40536 USA; Department of Biostatistics, Virginia Commonwealth University, Richmond, VA, 23298 USA

**Author notes:** Authors contributed equally to this work (Yelena Chernyavskaya, Xiaofei Zhang). Corresponding Authors: Jinze Liu, Jessica Blackburn.

**Keywords:** nanopore sequencing, zebrafish genome, reference assembly, transposons

## Abstract

Nanopore sequencing technology has revolutionized the field of genome biology with its ability to generate extra-long reads that can resolve regions of the genome that were previously inaccessible to short-read sequencing platforms. Although long-read sequencing has been used to resolve several vertebrate genomes, a nanopore-based zebrafish assembly has not yet been released. Over 50% of the zebrafish genome consists of difficult to map, highly repetitive, low complexity elements that pose inherent problems for short-read sequencers and assemblers. We used nanopore sequencing to improve upon and resolve the issues plaguing the current zebrafish reference assembly (GRCz11). Our long-read assembly improved the current resolution of the reference genome by identifying 1,697 novel insertions and deletions over 1Kb in length and placing 106 previously unlocalized scaffolds. We also discovered additional sites of retrotransposon integration previously unreported in GRCz11 and observed their expression in adult zebrafish under physiologic conditions, implying they have active mobility in the zebrafish genome and contribute to the ever-changing genomic landscape.

## Introduction

The relevance of model organisms to their human counterparts is strengthened by a high-quality reference genome. Complete genomic data allows for accurate evaluation of gene regulation, identification of mutations in disease states, assessment of evolutionarily conserved functional elements, and most importantly, permits manipulation of genetic sequence to create useful tools to study human diseases. However, most reference genomes contain regions of poor coverage, or gaps, that cannot be resolved with standard next generation sequencing (NGS) due to their short read size of 300 bases or less (Besser et al. 2018). Consequently, long-read sequencing technologies, such as Pacific Biosystems (PacBio) and Oxford nanopore sequencing, have emerged as a means to generate reads extending beyond 100Kbp, which are capable of spanning extended repeat regions to fill in genomic gaps.

The zebrafish has been used to study embryonic development since the 1960s (Anderson and Battle 1967; Weis 1968) but its more recent strength as a cancer and disease model has dictated the need for an accurate genomic assembly. Over 70% of genes associated with human diseases have a functional ortholog in zebrafish, and most human cancers can be engineered in fish by perturbing the counterpart orthologous genes (Santoriello and Zon 2012; Hason and Bartůněk 2019). Thus, having a quality reference genome is indispensable for molecular genetics in the zebrafish system.

However, several factors of the zebrafish genome complicate current assembly methods. First, the teleost genome has undergone multiple genome duplications, the most recent of which occurred after the divergence of the ray- and lobe-finned fishes more than 300 million years ago (Amores et al. 1998). Duplicate genes may exhibit redundance, dosage dependency, or other functions that are difficult to predict (Amores et al. 1998; Taylor et al. 2003; Espinosa-Cantú et al. 2015). Additionally, many of the duplicate regions exist on different chromosomes from one another or in a state where their identification, annotation, and mapping is difficult due to increased sequence divergence or existence on unlocalized contigs (Taylor et al. 2003). The second obstacle to assembling a high-quality reference for zebrafish is the excessive number of repeat regions present in the *Danio rerio* genome. A comprehensive study by Chopin et al., compared 23 vertebrate genomes and found that >50% of the zebrafish genome was represented by transposable elements (TEs) and repeats, higher than any other species examined, including human and mouse (Chalopin et al. 2015). These repeats can extend several mega base pairs and pose a formidable challenge to accurate assembly of the zebrafish genome, since sequencing-by-synthesis technologies cannot generate reads long enough to span these regions.

Recently, nanopore sequencing was used to access mobility of a 6 kilo base LINE-1 element in the human genome relative to its methylation status (Ewing et al. 2020). Although old TEs accumulate enough sequence diversity to be distinct from one another, young, mobile TEs are typically identical to their source element and cannot be resolved with short-read sequencing (Lanciano and Cristofari 2020). Nanopore sequencing overcomes the size constraints imposed by NGS, since native genomic DNA of any length can be fed through and “read” by each nanopore without the need for synthesis reactions (Jain et al. 2016). This allows for sequencing across repeat regions such as telomeres, centromeres and TEs (Miga et al. 2020; Jain et al. 2018; Ewing et al. 2020). Extended read length, however, is offset by lower base-pair accuracy, so most assemblies generated this way use supplementary short-read sequencing or increased depth to overcome this issue (Tyson et al. 2018; Miga et al. 2020).

According to the Genome Reference Consortium, the current zebrafish reference genome (GRCz11) contains 1,448 unresolved gaps, spanning across all 25 chromosomes, and 967 extrachromosomal unplaced contigs. Many of these regions are large enough to contain genes. However, because they lack concrete chromosomal locations their regulation remains a mystery since it’s impossible to know which cis-(promoters) or trans-(enhancers) acting elements govern their expression. Additionally, these statistics apply only to known issues with the current assembly and do not include errors that have yet to be defined. Recently, Yang et al. reported long-read sequencing of the zebrafish chromosome 4, demonstrating the capability of nanopore technology for filling in gaps resolving such issues (Yang et al. 2020). However, a genome-wide, comprehensive assembly has not been published. Here we report our findings and contributions from resequencing the zebrafish genome using a hybrid assembly of long-read nanopore sequencing and Illumina short-reads and demonstrate the ease and ubiquitous application of this recent sequencing platform in resolving difficult to map regions and genomic gaps. In addition to resolving the placement of formally unlocalized contigs and identifying new sequence indels, we’ve discovered novel retro-transposon insertion sites previously unreported in the reference assembly that contribute to genetic heterogeneity between different zebrafish model strains.

## Results

### Long-reads sequence across difficult genomic regions

According to the Genome Research Consortium, the major fraction of the 1,630 assembly issues within the zebrafish reference genome are gaps – ranging from a few thousand to several hundred thousand bases in length (Fig. 1). To span such large regions with substantial overlap, reads would need to approach gap length. To generate long reads essential for spanning these gaps and challenging repetitive genomic regions we tested two methods for purifying high-molecular weight genomic DNA. Since Tübingen served as the strain for the current reference genome we used a pool of muscle tissue from 4 Tübingen derived Sanger AB Tübingen (SAT) individuals for all library preparations. The first library (L180) was created with a standard in-house DNA extraction buffer and the second (L182) using the Nanobind Tissue Big DNA Kit (Supplemental Fig. 1)(Westerfield 2007). Kit extracted DNA produced consistently longer reads (N50 = 27.8Kbp) than the in-house method (N50 = 14.5Kbp) and was used for all subsequent library preps (Table 1; Fig. 2A; Supplemental Fig. 1). Sequencing was split across 6 different libraries, generating a total of 36.9 Gbp of sequence data. Although average read length was ∼ 15kb, a majority of the bases that were sequenced came from reads 20-150Kb in length, with the longest read spanning 464,751bp in L187 (Fig. 2A).

**Figure 1.**
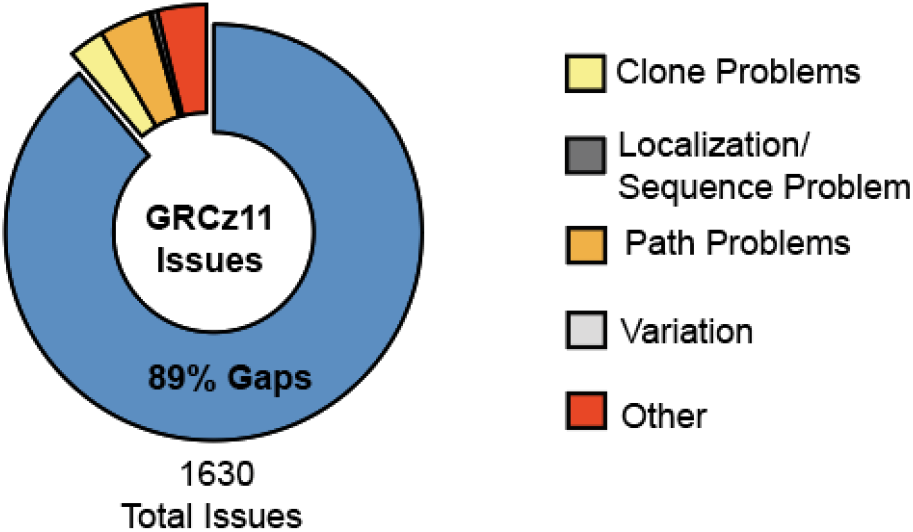
Curated current assembly issues with zebrafish reference genome GRCz11 as reported by the Genome Research Consortium

**Table 1:** Summary of Nanopore Sequencing Read Data for *D*.*rerio* SAT strain

| Library | Mean Read Length (bp) | N50 (bp) | Total Reads | Total Bases | Mean Alignment |
| --- | --- | --- | --- | --- | --- |
| L180 | 7,274 | 14,573 | 256,609 | 1.86676e+09 | 92.40% |
| L182 | 18,018 | 27,257 | 233,502 | 4.20726e+09 | 96.63% |
| L187 | 16,375 | 30,367 | 282,760 | 4.63028e+09 | 92.15% |
| L191 | 12,699 | 23,692 | 680,952 | 8.64755e+09 | 88.41% |
| L194 | 14,996 | 27,882 | 532,926 | 7.99194e+09 | 91.49% |
| L195 | 16,730 | 29,200 | 699,582 | 11.70513e+09 | 94.79% |

**Figure 2.**
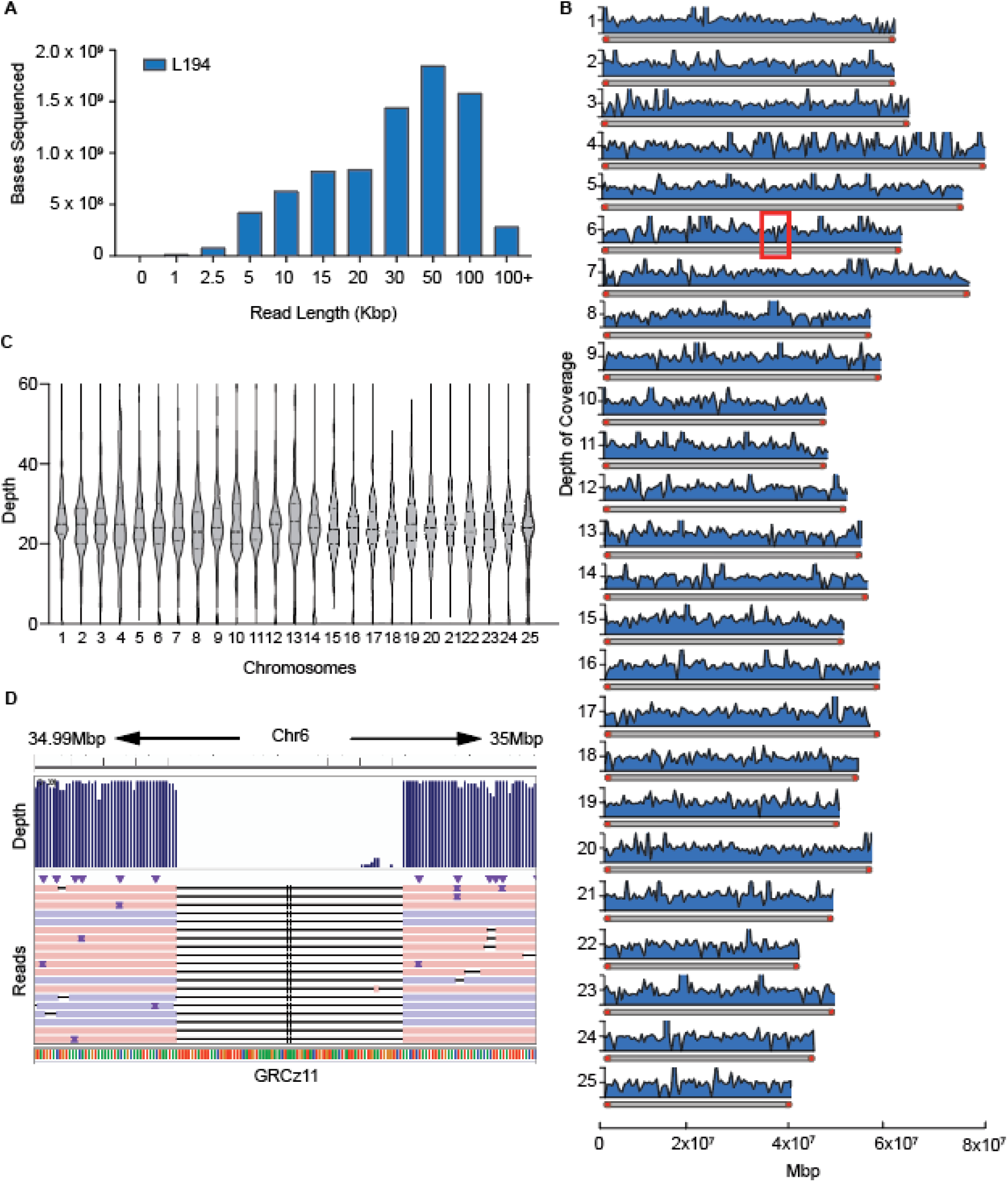
Long-read library run metrics. (A) Distribution of read lengths from one representative library relative to number of bases sequenced within that library. (B) Histogram of read depth and coverage across individual chromosomes at 50Kbp intervals. Chromosomes are depicted on the y-axis with maximum depth cut off at 50X. Telomeres (red caps) extend the first 20Kbp into each chromosome. Red box on Chr 6 emphasizes a region of low coverage. (C) Cumulative average depth across all chromosomes of long-read assembly. (D) Magnification of low coverage region depicted in B (red box) to show continuous nanopore reads spanning across the zero-coverage section of GRCz11.

Average sequencing coverage across the genome is used to assess general sequencing quality. However, this metric does not address the variability in coverage arising from sequencing across difficult DNA template. Such regions may not be adequately covered by reads, a factor that would be missed when averaging coverage across the entire genome. We determined how long reads generated by nanopore sequencing were distributed across the chromosomes and whether they were able to span notoriously difficult to sequence regions. Generally, reads were evenly distributed across all chromosomes – without over or under representation of any particular region – at an average depth of ∼30X (Fig. 2B-C). Next, we inspected how depth and coverage were represented at the terminal ends of zebrafish chromosomes. Since telomeres consist of repeat regions, it is inherently difficult to align short reads to them, resulting in a loss of information and accuracy at these important genomic locations (Galati, Micheli, and Cacchione 2013). Zebrafish telomeres are reported to extend 16-20Kbp into the chromosomes (Anchelin et al. 2011; Lund et al. 2009). Read depth at telomeres was slightly less than what was observed for the whole chromosomes, 24.3X for the left telomere and 28.9X for the right telomere respectively, but more than sufficient for long-read genome assembly (Supplemental Fig 2). The difference in depth between telomeres and intrachromosomal regions can be attributed to reduced number of sampling points at these locations.

Occasionally, we encountered areas of low sequence depth that justified further investigation. One such representative region exists at 35Mbp on Chr 6 (Fig. 2B, box on chr 6). Closer inspection of nanopore sequencing aligned to Chr 6 showed that all reads in that region were missing a 70bp section of sequence that is present in GRCz11 yet aligned accurately in every case on each side flanking the 70bp (Figure 2D). The presence of continuous, well aligned reads spanning both sides of the “low-coverage” region can be explained as an error in the original placement of that sequence in the reference genome and not an issue with the long-read assembly.

### Pipeline optimization for long-read genome assembly

To assemble the zebrafish genome *de novo* we compared two assembler tools, previously used for de novo assembly of large vertebrate genomes (Koren et al. 2017; Li 2016). Canu, originally developed for Pacbio, is an all-in-one package that overlaps, error-corrects and assembles long, noisy reads into contigs (Koren et al. 2017). Miniasm, on the other hand, requires a separate preceding overlap step and lacks built-in error correction, however its processing time is extremely short, which is an important factor to consider when dealing with large eukaryotic genomes (Li 2016). In addition, since nanopore sequencing is only ∼90% accurate, we opted for a hybrid assembly, incorporating several polishing steps using Illumina generated paired-end reads (McNaughton et al. 2019). Assembler statistics are summarized in Table 2.

**Table 2:** Summary statistics using two different assemblers relative to GRCz11

| Name | Polishing | Coverage | Total Contigs | Largest Contig | NG50 | Total Length | Run Time (d:h:m:s) |
| --- | --- | --- | --- | --- | --- | --- | --- |
| Canu | none | 90.8% | 3654 | 10,261,938 | 1,359,573 | 1,422,706,407 | 42:20:42:08 |
| Canu_RP | Racon + Pilon | 91.4% | 3523 | 10,335,318 | 1,383,746 | 1,432,333,057 | 43:14:47:52 |
| Miniasm | none | 88% | 1118 | 24,326,764 | 3,068,968 | 1,396,816,903 | 15:31:18 |
| Miniasm_RP | Racon + Pilon | 91.6% | 1118 | 24,721,058 | 3,165,400 | 1,417,315,502 | 1:01:31:41 |

As expected of assemblers with built in error-correction, Canu generated the largest assembly (1.42Gbp) with the highest coverage across the GRCz11 reference genome (90.8%) while Miniasm produced 1.39Gpb of sequence at 88% coverage (Table 2). However, correcting for base-pair errors with polishing packages (Racon and Pilon) reduced the variability in length and coverage between both assemblies. Although Canu has been commonly used for assembly of large genomes (Miga et al. 2020; Jain et al. 2018) we found that Miniasm surpassed it in genome coverage, contig length and NG50 (Table 2). When comparing assembly output in terms of contig lengths and numbers Miniasm_RP assembly covered the genome in only 1,118 contigs with the largest contig spanning an impressive 24.7Mbp and an NG50 of 3.16Mbp (Figure 3 and Table 2). In addition, Miniasm required a mere day to generate the assembly while Canu processing lasted almost a month and a half. Due to overall better performance, we chose the Miniasm generated and error-corrected assembly, hereafter referred to as ZF1, for all downstream analyses.

**Figure 3.**
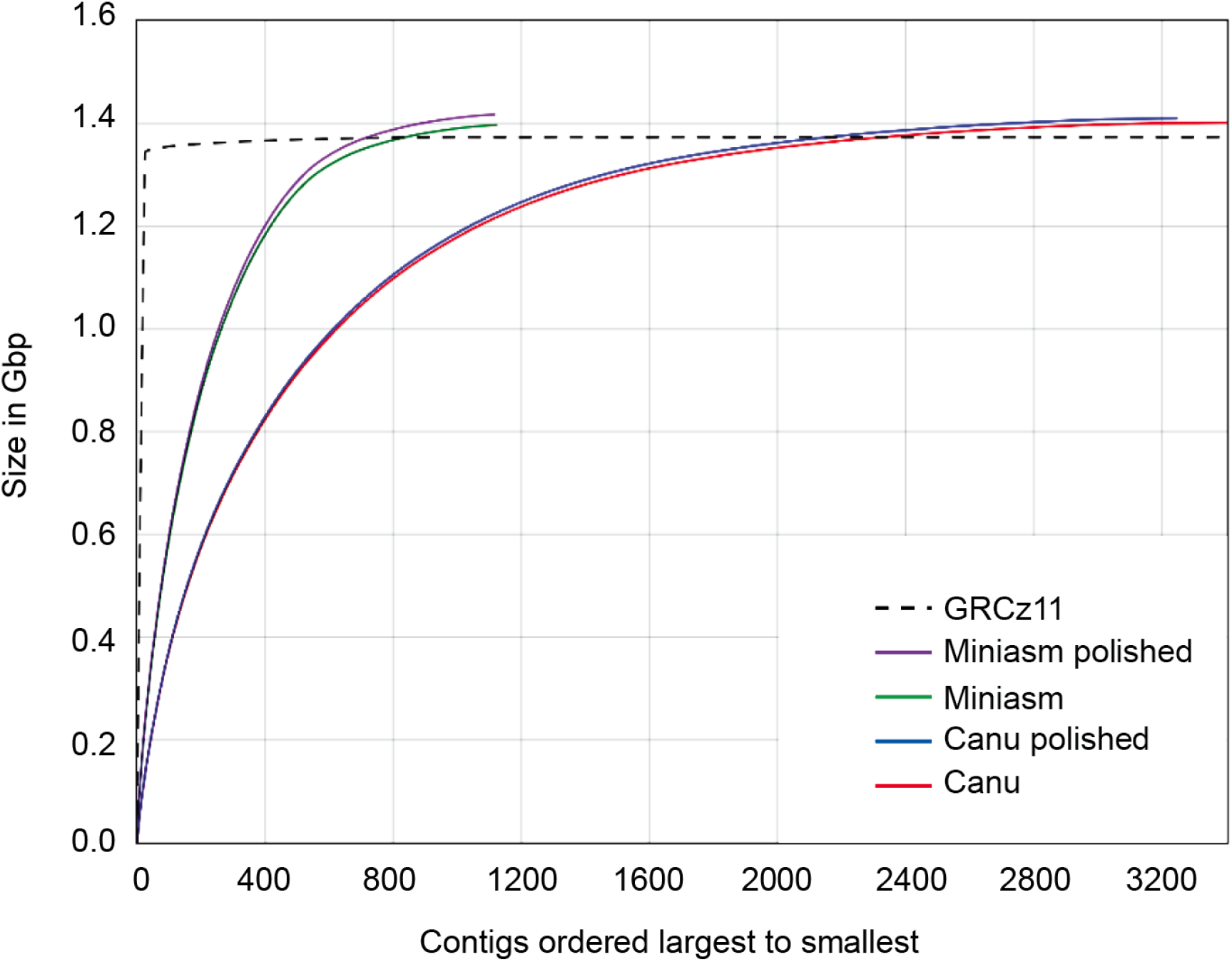
Comparison of assembly output versus number of contigs generated when using Canu and Miniasm with and without polishing steps.

### ZF1 assembly shows novel sequence placement

To assess the accuracy of our assembler pipeline we generated an association plot which diagrammatically depicts the alignment of two genomes. A strong association is represented by a solid diagonal line between the two axes. A comparison between our generated assembly and the reference genome showed a solid green line of contigs from ZF1 aligned to GRCz11 (Fig. 4A). However, there were key differences and variations, as indicated by small segments of alignment falling away from the diagonal line, between the assembly generated with long reads and the reference genome. For example, we identified a multitude of translocations and one large, 8.5 Mbp inversion, residing on Chr 2 (Fig. 4B). This inversion covers over 14% of Chr 2, contains 440 protein-coding transcripts and is large enough to span tandem associated domain (TAD) boundaries (Szabo, Bantignies, and Cavalli 2019; Pérez-Rico, Barillot, and Shkumatava 2020). We also focused on Chr 4 where we observed many small (<1Mbp) translocations present in ZF1 relative to GRCz11 (Supplemental Fig. 3). The reference sequence for Chr 4 is gene-poor and contains large gaps, making it one of the most poorly resolved zebrafish chromosomes. A similar pattern in translocation was reported by Yang et al., when they utilized long-read sequencing to map the *D*.*rerio* Chr 4, further supporting the validity of our long-read assembly (Yang et al. 2020).

**Figure 4.**
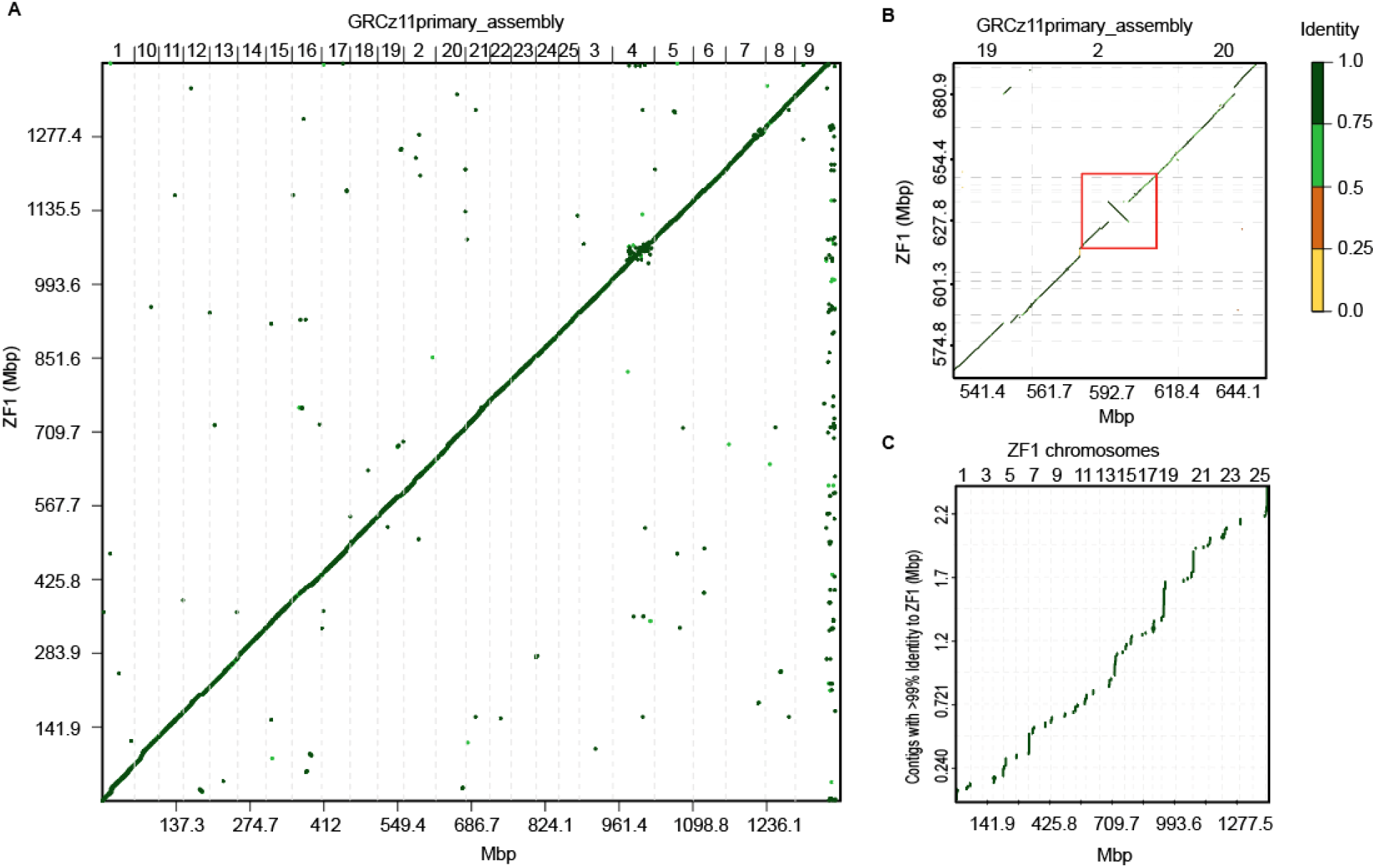
Association plots of similarities and differences between ZF1 assembly and GRCz11 primary assembly. (A) Entire *de novo* generated ZF1 assembly compared to GRCz11. Center, diagonal line marks strong association and alignments with shorter indels placed off-center of the diagonal (B) Magnified area on Chr 2 showing an 8.5Mbp inversion (red box) in ZF1 deviating from GRCz11. (C) Chromosomal placement of unlocalized contigs of GRCz11 bearing at least 99% similarity to ZF1. Color scale indicates percent similarity between alignments.

GRCz11 contains 967 unlocalized scaffolds, or sequences that are not localized to a position on any specific chromosome. Cumulatively, unlocalized scaffolds make up a total of 28.3 Mbp of unplaced genomic sequence in the zebrafish genome. Since the genome-to-genome association plot showed many small alignments off the diagonal we reasoned some of those could be newly placed unlocalized contigs from GRCz11. To determine whether our long-read assembly fully resolved the location of any of these we filtered out all scaffolds with a coverage below 99% within the ZF1 assembly. Scaffolds with lower coverages would only be partially placed in the new assembly. Placement of remaining unlocalized scaffolds showed that 106 had novel locations dispersed across all chromosomes of ZF1 assembly (Fig. 4C and Table S1). It is likely that the remaining unlocalized scaffolds suffer from low coverage at their junction points with the rest of the genome and could be assigned chromosomal location if sequencing depth was increased.

### Novel Chromosomal Indels in ZF1 contain LTR Transposons

Next, we focused on identifying and curating total novel insertions and deletions within the ZF1 assembly. Considering that nanopore based sequencing has a base-calling error rate of ∼10%, higher compared to more conventional methods, we set a 1,000bp threshold for all novel genomic elements identified, since insertions or deletions (indels) of that size are unlikely to be caused by assembly mistakes generated from base-calling errors (Wick, Judd, and Holt 2019). In total, we identified 1,049 insertions and 648 deletions of >1,000bp across the entire zebrafish genome (Fig. 5A). We found no correlation between indel frequency and chromosome size, suggesting that indels did not randomly increase in number with increasing chromosome length (Fig. 5B). Instead, indel frequency is probably a factor of sequence complexity since chromosomes harboring more repeat elements are more likely to have assembly issues. To determine if any deletions in ZF1 stemmed from mis-localized genomic sequence in GRCz11, we cross-referenced the deletions to the insertions with a minimum cutoff of 98% identity and 98% coverage. This assessment revealed that 93% (n = 603) of the original deletions identified in ZF1 had novel placements in other parts of the assembly (Table S2).

**Figure 5.**
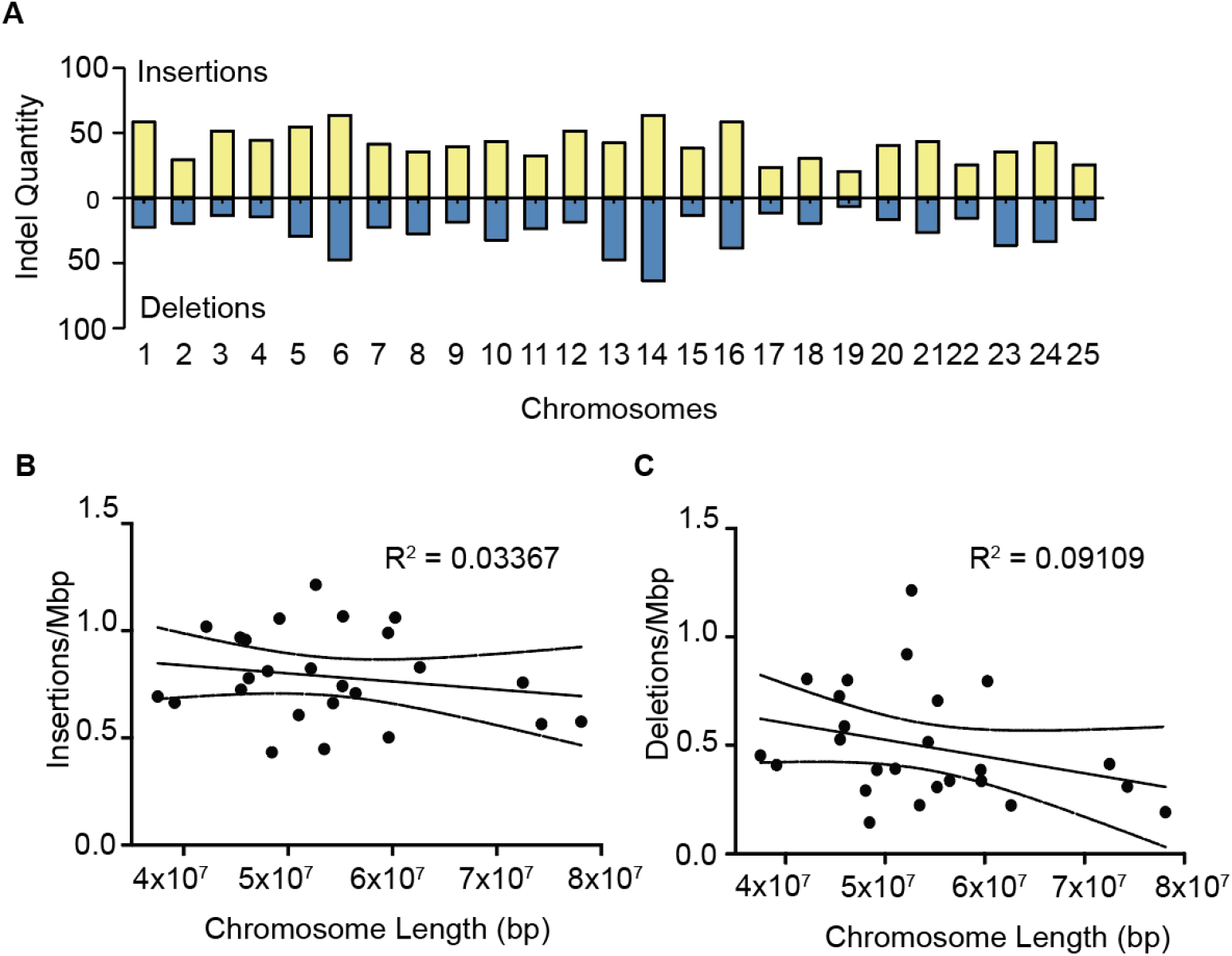
Novel indel distribution in ZF1 assembly. (A) Frequency of insertions (yellow) and deletions (blue) identified in ZF1 assembly across all chromosomes. (B-C) Dot plots showing lack of correlation between indel frequency and chromosome length.

Insertions of >1,000bp are large enough to contain genetic elements whose regulation is dictated by their genomic location. We mined the 1,049 insertions with gene prediction software to locate potentially new genes. Geneid detected 23 protein coding genes, all belonging to the LTR Retrotransposon family (Fig. 6A). Since repetitive elements are often difficult to map, we expected most of these LTR retrotransposons to also be present in the deletion dataset, indicative of their original misplacement in the reference genome. We chose 4 representative LTR retrotransposon indels from the 23 candidates for interrogation their original genomic coordinates in GRCz11. Considering that specific LTR retrotransposons can occur numerous times in the genome, we investigated every occurrence as a potential source of the indel. To reduce the chance of mistaking one LTR retrotransposon species for another due to high sequence similarity, we used a minimum identity cutoff of 99%. The 4 indel LTR retrotransposons from ZF1 were BLASTed against GRCz11 to obtain their original location and then that region was compared against ZF1 for the presence of the LTR retrotransposon of interest. Three of the 4 LTR retrotransposons interrogated retained all their genomic locations from GRCz11 in ZF1 (Table S3), while Gypsy52-I_DR was missing in 2 of its 5 genomic coordinates in ZF1, possibly due to original assembly errors. These data indicate that strain-specific differences exist within the zebrafish that deviate from the published reference genome. Since assembly errors in GRCz11 could not explain all the novel insertions of the interrogated transposons we presumed their integrations might be due to activity in the genome.

**Figure 6.**
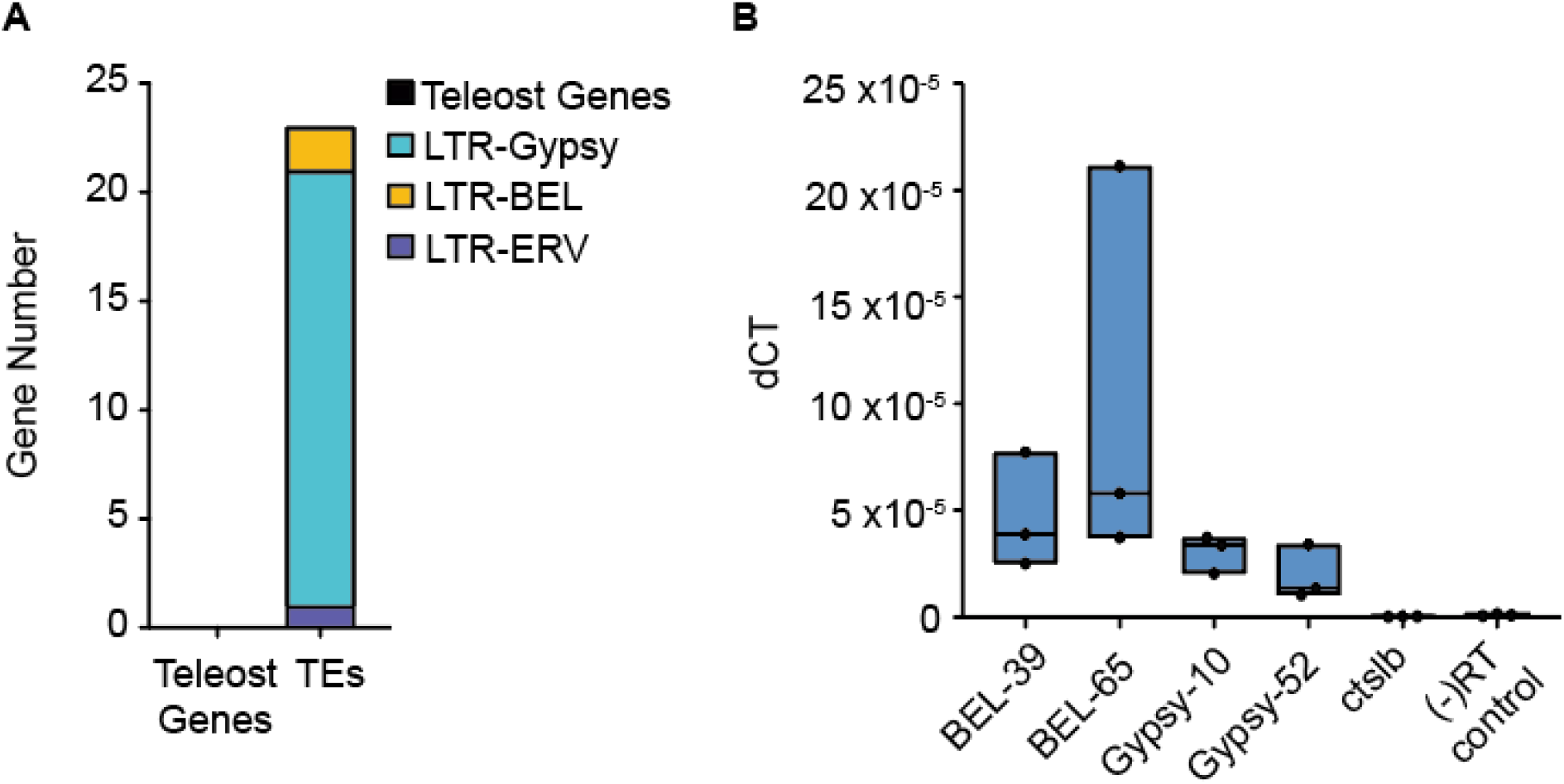
Identification of active retrotransposons in ZF1 assembly. (A) Results of gene prediction software reveals 23 novel insertions of LTR retrotransposons in *de novo* assembly. (B) Expression by RT-qPCR of select retrotransposons compared to *ctslb*, which is silenced, and a negative control amplified with primers meant to pick up genomic DNA contamination.

Although it is commonly believed that most transposable elements are silenced by the host cell, reported instances exist showing their expression and activity is necessary under physiological conditions for regulation of gene expression and as functional components of nuclear architecture in early embryonic development (Johnson and Guigó 2014; Todd et al. 2019; Percharde et al. 2018). LTR retrotransposon mobility depends on the presence of expressed mRNA which is reverse-transcribed and re-inserted into new sites in the genome (Wicker et al. 2007). In this manner their activity manifests as novel genomic integrations while retaining the placement of their original copies. To investigate LTR retrotransposon activity, we monitored the expression levels of the 4 indel LTRs in 3-week-old zebrafish and compared these to the mRNA abundance of cathepsin Lb (*ctslb*), which is silenced post hatching (Figure 6B). As expected, expression level of *ctslb* was almost undetectable. Primers designed to amplify genomic DNA in the absence of reverse transcription produced low signal, indicating that samples have very little genomic DNA contamination. LTR expression, however, was present above that of the silenced gene or possible genomic contamination, confirming their expression in the host cell.

## Discussion

Since its advent, applications of nanopore sequencing for genome reconstructions have risen exponentially. Although several long-read vertebrate reference genomes now exist, including human and *C. elegans*, a complete genome assembly of zebrafish had not been released and the current GRCz11 reference is still replete with a multitude of assembly issues (Fig. 1) (Yoshimura et al. 2019; Jain et al. 2018). Long-read sequencing of the human genome was used to resolve telomeres and centromeric repeats, a difficult task for earlier sequencing technologies, while sequencing of *C. elegans* strain VC2010 led to the discovery of strain specific differences including 53 new genes and the addition of 1.8 Mbp of novel sequence (Miga et al. 2020; Yoshimura et al. 2019). Previous challenges in assembling the zebrafish genome stemmed partly from technological limitations of short-read sequencing but also from the complexity of deciphering the throng of repetitive elements that comprise more than 50% of the entire genomic landscape in this species (Chalopin et al. 2015). Additionally, an overabundance of repetitive sequence can contribute to PCR artifacts during library preparation, which can further confound mapping and assembly. Thus, we set out to resolve the current reference genome issues plaguing GRCz11 by using native DNA, long-read sequencing to create a more accurate zebrafish assembly.

To generate the most accurate assembly build, we compared 4 different pipelines utilizing the two assemblers currently used for long-read vertebrate genomes. Although the Canu generated assembly was slightly larger than its counterpart created with Miniasm, Miniasm outperformed Canu in several quality metrics such as NG50, total contig number and size. In addition, Miniasm required mere hours to complete the assembly process compared to the incredible CPU requirement of more than 40 days for Canu. In total, our ZF1 assembly added 43.86Mbp of sequence to the zebrafish genome, equivalent to the size of an entire chromosome, and imparted chromosomal coordinates to 107 scaffolds previously unlocalized in GRCz11. We also identified a large 8Mbp inversion on Chr 2, which holds potential biological significance since its size is large enough to encompass multiple regulatory regions such as topologically associated domains (TAD), (Dixon et al. 2012; Pérez-Rico, Barillot, and Shkumatava 2020). TADs are structural chromosomal domains that maintain preferential intra-domain interactions and are subject to gene regulation based on their location and placement relative to other long-range enhancers, thus a gene might be regulated differently whether it’s in one TAD or another (Szabo, Bantignies, and Cavalli 2019). The 8Mbp inversion completely re-organizes the placement of hundreds of genes, whose regulation would be subject to change as well based on their updated genomic coordinates.

Similarly, we also investigated placement errors in the GRCz11 relative to ZF1. we identified a total of 1,697 insertions and deletions greater than 1Kb. Most (608/648) deletions were also represented in the insertions group, suggesting they were misplaced in the original reference genome. Further examination of indels identified 23 LTR retrotransposon genes present within the insertions. This finding was not surprising since transposable elements are so widespread in the zebrafish genome; however, LTR retrotransposons encompass only 10% of all transposable elements, with DNA transposons representing 80% of the group (Chalopin et al. 2015). The probability of randomly encountering an LTR retrotransposon within the insertions would therefore be low relative to DNA transposons or other repeats. In addition, we found that most copies of the newly identified LTR retrotransposons were retained between GRCz11 and ZF1, suggesting that the insertions were not due to previously misplaced LTR retrotransposable elements but instead indicative of additional insertion mechanisms beyond random error, such as the reverse transcription/reintegration method utilized by active retrotransposons. Although direct assessment of TE mobility was beyond the scope of this study, we did assess the expression of 4 select retrotransposons and found them to be expressed above the level of repressed genes, suggesting their activity in the genome, at least at the transcriptional level. Transposons can be and are active throughout important biological and developmental events, such as immune priming, and domestication of retrotransposons is one mechanism by which genes form (Chuong, Elde, and Feschotte 2017; Chernyavskaya et al. 2017; Wang, Tracy, and Zhang 2020; Kapitonov and Koonin 2015; Zhang et al. 2019). Gypsy, for example is documented to be mobile and infectious in Drosophila, actively remodeling the genomic and regulatory landscape in this organism (Kim et al. 1994; Nefedova and Kim 2017; Wang, Tracy, and Zhang 2020). Although a genome wide assessment of transposon mobility has not been carried out for zebrafish, our data strongly suggests that retrotransposons are active in the genome of adult *Danio rerio*. Thus, gene regulation within the genome should be considered dynamic and strain specific in light of retrotransposon contribution, which is ongoing and ever present.

## Methods

### DNA extraction and library preparation

To generate high-molecular weight (HMW) genomic DNA 4 adult Sanger AB Tubingen (SAT) fish were sacrificed by tricaine overdose. Tail muscle tissue from all 4 was pooled and flash frozen in 25mg aliquots. DNA extraction for the 1^st^ library was carried out using a house-made extraction buffer (10mM Tris pH 8.2, 10mM EDTA, 200mM NaCL, 0.5% SDS, and 0.2mg/ul Proteinase K) and the Westerfield DNA extraction protocol (Westerfield 2007). All subsequent libraries were generated with DNA extracted using the Nanobind Tissue Big DNA Kit (Circulomics NB-900-701-01) using their Standard TissueRuptor Protocol – HMW. Following extraction DNA was allowed to rest 24-48hr to solubilize into the solution. Solubilized DNA was size selected with SRE Short Read Eliminator Kit (Circulomics SS-100-101-01) according to manufacturer’s protocol and 1.5µg was used as input for library prep. Six libraries were generated using the Oxford Nanopore 1D Genomic DNA by Ligation Sequencing Kit (SQK-LSK109) according to the protocol provided except for the following optimizations for HMW DNA. The End Prep/Repair step was increased to 60min, and the Adapter Ligation incubation was carried out for 10hrs. 400-600ng of each prepared library was loaded in 75µl volume onto flow cells and run for 24-30hrs, until flow cell extinction, for an average N50 of 27.2Kb. An aliquot of the gDNA used for nanopore library prep was also used for paired-end Illumina whole-genome sequencing carried out by GeneWiz.

### Assembly Pipeline

Fast5 data was base-called using Guppy and all mapping was performed with Minimap2 (v2.16). The Samtools (v1.10) was used in index generating, alignment file sorting and alignment statistics calculations. Assemblies were generated using two pipelines. The first used Canu (v1.9) and the following source code: canu -d ../Canu -p ZF1 genomeSize=1.4g useGrid=false -nanopore-raw ../FASTQ/ZF1.fastq. The second used Minimap2 to first generate the pairwise mapping (PAF) file: minimap2 -x ava-ont - r 10000 -t 16 ../FASTQ/ZF1.fastq ../FASTQ/ZF1.fastq> ../MINI_OUT/ZF1_overlap.paf. This was used as input for Miniasm (v0.3) to create the assembly: miniasm -f ../FASTQ/ZF1.fastq ../MINI_OUT/ZF1_overlap.paf > ../MINI_OUT/ZF1.gfa. The awk was used to write the assembly file: awk ‘$1 */S/ {print “>“$2”\n”$3}’ ../MINI_OUT/ZF1.gfa > ../MINI_OUT/ZF1.fasta

Polishing was performed in two ways. First, the pairwise mapping format files of the unpolished assembly and the raw long reads were generated using Minimap2: minimap2 -t 16 ../MINI_OUT/ZF1_MM.fasta ../FASTQ/ZF1.fastq > ../MINI_OUT/ZF1_overlap_for_polishing.paf, followed by Racon (v1.4.13) to polish the unpolished assemblies using the raw long reads. Next, short-read polishing using Pilon (v1.23) was performed using Illumina whole-genome sequencing data. The alignment files of raw reads to the assembly were first generated using bwa (v0.7.17) and indexed using Samtools (v1.10). Then the Pilon (v1.23) was used to polish the assembly using the short reads alignment: pilon -Xmx160G --genome ./FASTA/ZF1_MM_R_lr.fasta --fix all --changes --bam ./BAM/ZF1_MM_R_lr_sr_mapping.sorted.bam --threads 32 –output ./pilon_canu/pilon_round1 | tee ./pilon_canu/round1.pilon.

### Variant calling, genetic element identification and association plots

The paftools.js in Minimap2 (v2.16-r922) was used to call variants from the generated assembly against the reference. minimap2 -cx asm5 –cs

./ZF_Ref/Danio_rerio.GRCz11.dna.primary_assembly.fa

./Assemblies/ZF1_MM_R_lr_R_sr.fasta \ | sort -k6,6 -k8,8n \ | paftools.js call -f

./ZF_Ref/Danio_rerio.GRCz11.dna.primary_assembly.fa - >

./VCF/ZF1_MM_R_lr_R_sr.vcf. From the generated VCF file, the indels with size larger than or equal to 1000 bases were selected by checking the sequencing lengths of the ‘REF’ and ‘ALT’ column for each variant. The involved sequences were written in a FASTA file.

Genetic elements within the insertions from the VCF calling were predicted using Geneid (v1.4), using the human parameter file ‘human3iso.param’ which can be used for vertebrate genomes and the following compands: geneid –XP

/home/xzh289/Tools/geneid/param/human3iso.param

./1000bp_insertion/ZF1_MM_R_lr_P_sr_1000bp_insertion.fasta>

ZF1_MM_R_lr_P_sr_1000bp_insertion.extend.gff. Newly discovered genetic elements were than BLASTed to confirm their identify or conserved motifs.

Association dot plots comparing the ZF1 assembly to GRCz11 reference or to GRCz11 unlocalized contigs bearing >99% identity in ZF1 were carried out using the web-based version of D-Genies and .paf files previously generated by Minimap2 (v2.16-r922) (Cabanettes and Klopp 2018).

### Retro-transposon locations and expression

To determine if the LTR retrotransposons identified with Geneid (v1.4) maintained their original GRCz11 genomic locations in ZF1 we mined the alignment data of ZF1 assembly to GRCz11 reference at those locations where the LTR retrotransposons of interest were shown to exist (Supplement Table 1). To assess expression of the four retrotransposons, RNA was extracted from 3 week old SAT fish using TRIzol™ Reagent (Fisher Scientific) according to manufacturer’s protocol and all residual DNA was removed using DNA-free™ DNA Removal Kit (Life Technologies). Real-time quantitative PCR (RT-qPCR) primers were designed to span 120-170bp region of each LTR-RT. As a control for monitoring transcript abundance of genes that should not be expressed in adult zebrafish, we also included primers for cathepsin Lb (*ctslb*), a peptidase expressed in the hatching gland during early larval development.

## Data Access

This Whole Genome Shotgun project has been deposited at DDBJ/ENA/GenBank under the accession JAIHOL000000000. The version described in this paper is version JAIHOL010000000.

## Acknowledgements

We would like to thank Jeremiah Smith and Dylan Rivas for advice regarding data processing. Funding in support of this project was provided by the National Institutes of Health DP2CA228043 (JSB) and the Kentucky Pediatric Cancer Research Trust Foundation (JSB). This research was also supported by the Biostatistics and Bioinformatics Shared Resource Facility of the University of Kentucky Markey Cancer Center (P30CA177558) and the Virginia Commonwealth University Massey Cancer Center Bioinformatics Core (P30CA016059-31S1).

