## Supplemental Figures for "Long-Read Sequencing of the Zebrafish Genome Reorganizes Genomic Architecture"

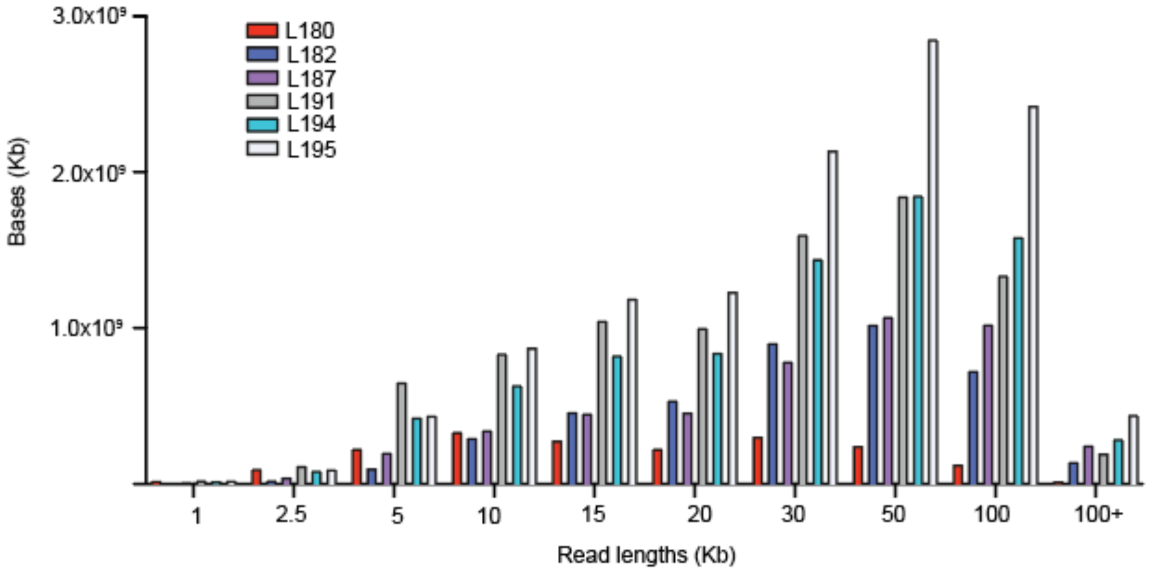


**Supp. Figure 1**. Read length distribution and sequenced bases generated by each group across all libraries used in assembly generation.

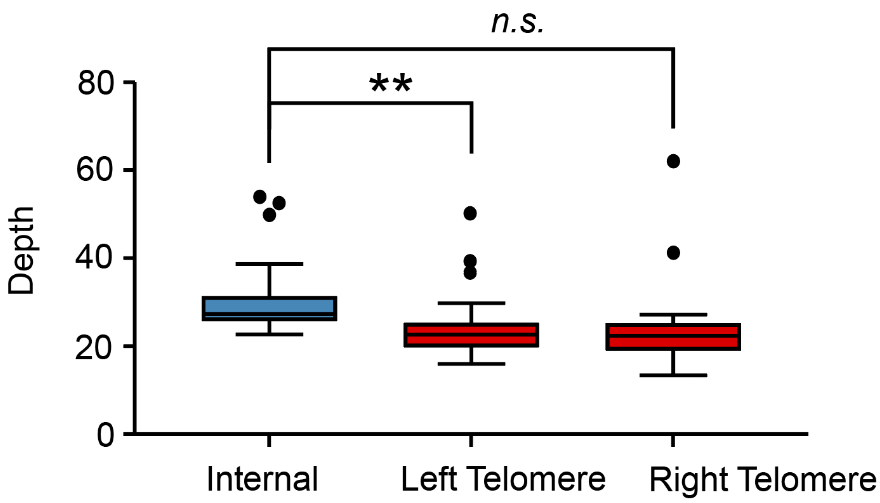


**Supp. Figure 2**. Tukey box and whiskers plot of average depth at the telomeric regions of all chromosomes in zebrafish genome. Significance between telomeric and intra-chromosomal depth was calculated by t-test, p-value 0.0097 (**), *n.s.* is no significance.

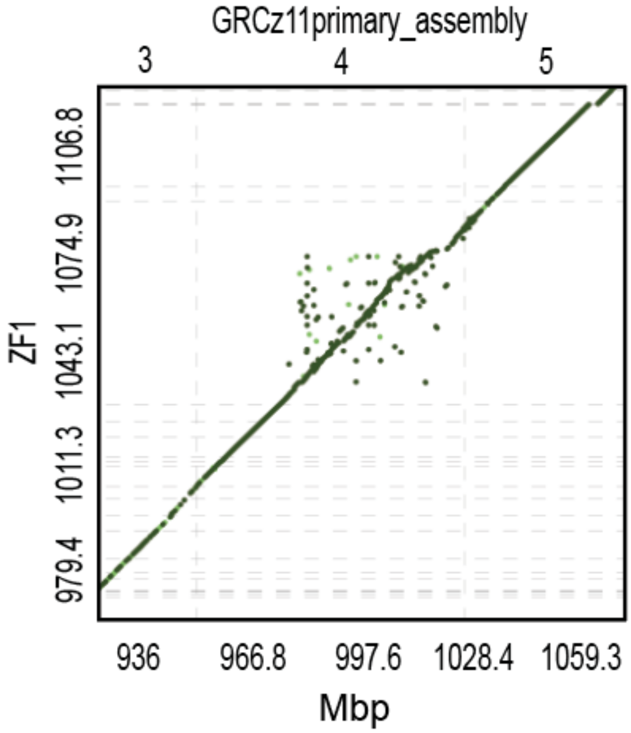


**Supp. Figure 3**. Association plot of Chr 4 in ZF1 and GRCz11 assemblies illustrating many small sequence differences between the two builds.
